# Post-fire *Quercus alba* fitness in a stressed plant community

**DOI:** 10.1101/189829

**Authors:** Kevin Milla

## Abstract

Prescribed burns are widely used for managing North American deciduous forests due to their ability to positively affect plant community structure and composition. This study examines the effects of neighboring herbaceous plants on the recruitment of *Quercus alba* (white oak) seedlings in fire-managed parts of Shawnee National Forest (Illinois, USA). Herbs were clipped to induce plant community stress and relative growth rates (RGRs) of planted white oak seedlings were assessed to determine if a competitive or facilitative dynamic is present. In addition to RGR, we observed the mycorrhizal network via fungal colonization in mesh bags to quantify belowground activity for our experimental plots. Our results supported fire’s positive effects on tree recruitment and herbaceous growth. Clipping combined with fire management decreased RGR. This finding suggests that a facilitative dynamic is at play and herbaceous neighbors help white oak seedlings persist due to protection from environmental stressors (p = 0.017). Soil moisture played a large role in promoting tree fitness on each of our sites. Lower hyphal biomass was observed in areas where herbs were clipped. We further speculate that the stress caused by clipping may have suspended or eliminated the need for mycorrhizae to form, possibly due to herb mortality. Knowing how herbs and trees interact will lead to purposeful forest community planning especially in fire-managed forests where herbs are likely to dominate post-prescribed burn.

## 1. Introduction

The idea that multiple species can comfortably coexist in any given space is an understated phenomenon. As more organisms start to populate a community, competition for resources becomes inevitable. The nature of ecological research demonstrates that competition for limiting resources happens in almost every ecosystem. In forests, the composition and distribution of trees and grasses are a product of their individual strategies to secure light, water, and other resources. If one species outcompetes another, it can lead to a variety of detriments for the less advantageous species, often leading to its demise. Consequently, post-successional patterns are easily predicted with the victors of competition more likely to achieve reproduction and fit to be successful at spreading offspring. Plant-to-plant competition is the interest of many studies aiming to predict productivity of a species or in creating guidelines for management and protection of ecosystems.

Facilitation, the opposite of competition, highlights the positive interactions between species in a community. The implied collapse of inferior competitors in a forest is juxtaposed by the thought that trees would benefit in the presence of neighboring plants, especially in its early lifespan. Younger trees are more susceptible to abiotic stress because seedlings have relatively less carbon available in their internal storage and far less access to deep soil water reserves compared to larger mature plants (Niinemets 2010). Previous studies have established that plant communities increase relative humidity, regulate temperature fluxes, and prevent over-drying of soil, all of which are classified as aboveground microclimatic changes (Holmgren et al. 1997, Classen et al. 2010, Montgomery et al. 2010). These changes are measured instantaneously within the O and A layer of the soil horizon and are averaged over short periods. Plant communities, therefore, utilize the humid microclimate to protect themselves from environmental stressors (Wright et al. 2014). Additionally, the persistence of trees and plants together over longer temporal scales can be characterized by their belowground mycorrhizal network.

The mycorrhizal network is meant to support a plant community in times of stress. The presence of ectomycorrhizae (EM) is a sign of root activity in most trees, and the presence of arbuscular mycorrhizae (AM) is a sign of root activity in most herbs (Wang and Qiu 2006). This complex relationship is characterized by the exchange of nutrients and water through the soil and into the roots of the plant. The hyphal network protects the plant in the event of prolonged biotic or abiotic stress (Barea and Pozo 2013). Likewise, according to the stress-gradient hypothesis, facilitation is more common in plant communities under high abiotic stress (Maestre et al. 2009). Our experiment will focus on the fitness of Quercus alba (white oak) trees surrounded by herbaceous plants in an oak-hickory forest. To explore aboveground and belowground changes that affect white oak fitness, we chose conditions and performed treatments that induce the effects of stress in a plant community.

We decided to study oak-hickory forests that have been fire-managed. Fire is a relatively common forest disturbance type that promotes species abundance, richness, diversity, and evenness in understory vegetation as well the subtraction of diseased and dead trees and litter (Brose and Van Lear 1999). Fire science has led to real-world management applications in at least million acres of forest in North America (Melvin 2015). In the absence of fire management, the reduced light availability for herbaceous plants and the accumulation of forest floor material may prohibit establishment of herbaceous vegetation and new oak generation. Light availability is moderately important for the establishment of oaks, which are intermediately shade tolerant (Moore 2002). In central and eastern mesic regions, prescribed burns are used to remove fast-growing competitors to the fire-tolerant oak (Knapp et al. 2015). The herbaceous matter is top-killed, but the competitors’ fast resprouting ability is responsible for the succession from a woody-dominated to an herbaceous-dominated understory after a fire (Peterson and Reich 2001). Low-intensity burns are utilized around the world for many land management purposes, but it is sparsely studied as to why resulting herbaceous-dominated understories benefit oak recruitment. We isolated the interactions of both fire and plant stress to evaluate if tree fitness relies on either competition or facilitation with neighboring herbs.

This experiment makes assumptions regarding broad generalizations about competition and facilitation and the initial conditions of both burned and unburned plots. Competition and facilitation can coexist in an ecosystem in varying degrees, and it is wrong to assume either behavior absent, or one solely responsible, due to present evidence of either competitive or facilitative behavior. From this point on, competitive behavior and facilitative behavior will be referred to as just competition or facilitation, without further mention that both may be found in our study areas under more involved methodologies. Furthermore, in our research sites, we assume initial AM abundance to be relatively smaller in fire-absent plots due to fire-managed plots exhibiting herbaceous-dominated understories. Nevertheless, only total hyphal biomass was measured, and we did not differentiate between EM or AM in this experiment.

Over two years, we manipulated the amount of herbaceous matter by clipping them from our research sites to simulate stress within burned and unburned plots. We also deployed mesh bags each year to measure hyphal biomass for every variation of treatment. Our hypotheses are as follows:

1. A low-intensity fire regime will increase the relative growth rate (RGR) of white oak seedlings in an eastern deciduous hardwood forest.
2. Removal of neighboring herbaceous plants by clipping in a fire-managed forest will increase RGR of white oak due to reduced competition for light and water.
3. Removal of neighboring herbaceous plants by clipping in a fire-managed forest will decrease RGR of white oak due to reduced protection from environmental stressors, demonstrating facilitation.
4. A mycorrhizal network of EM-AM fungi will increase as a result of induced herbaceous plant stress by clipping in an eastern deciduous hardwood forest.

Our data indicates that facilitation is present due to negative effects on white oak fitness resulting from induced stress on the neighboring plant community.

## 2. Methods

### 2.1. Study site

We conducted this study in Shawnee National Forest in southern Illinois, USA (37°26’N, 88°67’ W) from 2014 - 2015. The region is a western extension of the unglaciated Interior Low Plateau with a bedrock of Pennsylvanian and Mississippian sandstone and some interbedded limestone. The sites were located on south-facing slopes, where the bedrock is topped with a shallow layer of loess. The forest canopy species of these xeric sites are dominated by *Q. alba, Q. marilandica, Q. rubra, Q. stellata, Carya glabra, C. laciniosa, C. ovata* and *Juniperus virginiana*.

### 2.2. Experimental design

In 2014, we selected two sites under active prescribed burn management (burned ≥ 3x within the last 15 years by the US Forest Service), each paired with a nearby (≤ 1 km) unburned reference site. Unburned sites had no record of fire activity in the past three decades, according to United States Forest Service records dating back to 1980 (Scott Crist, pers. comm.). A paired block design containing an unbalanced distribution of four plots within each burned site and two plots within each unburned site was used to maximize our ability to assess herb removal treatment effects, given herb removals would be minimal in unburned plots during initial plot set-up.

Each plot (24 × 3 m) contained eight subplots (3 × 3 m), each randomly assigned to one of four treatments: +Seedling, -Herb, +Seedling / -Herb, and Control. This design ensured each treatment was replicated once per plot. Treatments were applied to a 2 × 2 m area in the center of each subplot to provide a 1 m buffer around treatments (Figure 1). For +Seedling subplots, ten bareroot *Quercus alba* (white oak) seedlings, sourced from the Illinois Department of Natural Resources, were planted in early May 2014 in a regularly spaced grid (total seedlings = 480). For -Herb subplots, all herbaceous vegetation was clipped at ground-level using hand-clippers. We chose to clip rather than pull vegetation to minimize soil disturbance. Clipping was conducted three times during each growing season, for a total of six times over the duration of the study. With the exception of the first clipping (see *Understory percent cover*), all clipped biomass was left in each subplot rather than removed. Woody species, including vines, were not removed (e.g. *Parthenocissus quinquefolia, Toxicodendron radicans, Rubus spp., Smilax spp.*, and *Vitis spp.*). Control subplots were not clipped.

**Figure 1:**
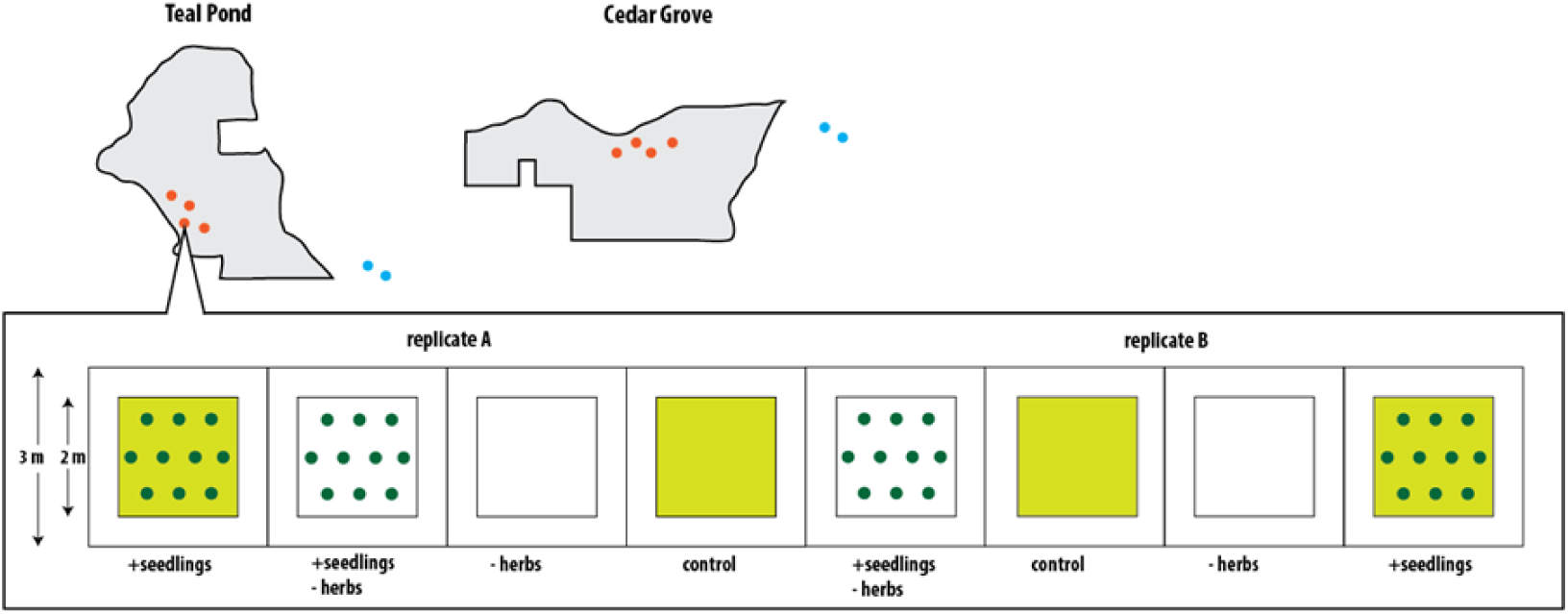
Map of study sites in Shawnee National Forest with burned (red dots) and unburned (blue dots) designations. Sample replicates are shown with all treatment variations: +Seedlings (green dots with yellow box), -Herbs (white box), +Seedlings / -Herbs (green dots with white box), Control (yellow box).

### 2.3. Soil parameters

In 2014 and 2015, bulk density, soil pH, and gravimetric soil water content (SWC) were measured in each subplot (n = 96). To determine SWC (%) and bulk density (g soil cm^−3^), we collected one 5 cm diameter soil core to 10 cm depth in July, sieved to 2mm and oven-dried (105°C) a 10 g subsample of the ≤ 2mm fraction. To determine soil pH, we collected two 2.5 cm diameter soil cores to 10 cm depth in July. The sample was sieved to 2mm, air dried and then measured for pH using a 2:1 ratio of deionized water to air-dried soil.

### 2.4. Understory percent Cover

To estimate the effect of clipping on percent cover of understory vegetation, we photographed the vegetation cover in each subplot before and after the first clipping treatment in late May 2014. Images were analyzed for total understory cover in Adobe Photoshop (Adobe Systems, Mountain View, California, USA) by estimating the percentage of pixels in each subplot that contained green vegetation. During the first clipping treatment in May 2014, all clipped biomass was collected, air dried and weighed in order to establish a relationship between total percent cover estimates and aboveground herbaceous biomass removed by clipping.

In June 2015, percent ground cover in each subplot was visually estimated into one of seven categories: 0%, 1-5%, 5-25%, 26-50%, 51-75%, 76-100% (Daubenmire 1959). Cover was estimated separately for each of five functional group: arbuscular mycorrhizal-associated tree species, ectomycorrhizal-associated tree species, shrubs or vines, forbs and graminoids. Bare ground, litter, and rocks cover were not estimated.

### 2.5. Light availability

We estimated understory light availability in July 2014 and 2015 using a LI-191 Line Quantum Sensor (LI-COR Biosciences, Lincoln, Nebraska, USA). Photosynthetically active radiation (PAR) was quantified approximately 0.5 m above each subplot and under full sun in canopy gaps. Light availability was estimated by dividing subplot PAR measurements by the PAR reading of the canopy gap (%). Readings were taken on cloudless days between 10:00 - 14:00.

To estimate canopy basal area, we measured the diameter at breast height (DBH) of all adult trees (≥ 10 cm DBH) in a 34 × 12 m (0.04 ha) area surrounding each 24 × 3 m plot. Measurements were taken in July, 2015. Individuals were identified to genus, with the exception of *Quercus spp*. were individuals were identified as either red (sect. *Erythrobalanus*) or white oaks (sect. *Lepidobalanus*).

### 2.6. Seedling relative growth rate and leaf area

We measured seedling basal diameter, height and number of leaves at the beginning and end of each growing season (30 May 2014 - 25 July 2014 and 15 May 2015 - 28 July 2015). To estimate the relationship between size measurements and aboveground biomass (AGB), we harvested *Q. alba* seedlings near the study plots, dried and weighed all aboveground components of each individual. For multi-stemmed individuals, AGB was calculated separately for each stem and then summed together.

AGB (g) was calculated using the following equation:

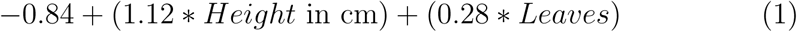

RGR was calculated using this equation:

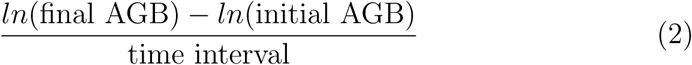

(Poorter and Lewis 1986).

To estimate leaf area, we collected three undamaged leaves from each individual after the final census, in late July. Each leaf was scanned and the area estimated (cm^2^) using Adobe Photoshop.

### 2.7. Hyphae production

In 2014, fungal in-growth bags were made from nylon mesh (53 *μ*m mesh size, 10 × 5 × 2 cm) by sealing edges using Amazing Goop All-Purpose Adhesive (Eclectic Products, Inc., Eugene, Oregon, USA). The size of the mesh allowed for colonization of fungal hyphae, without root penetration. After the bags were filled with sterilized coarse sand, the adhesive was left for 24 hours to cure.

Mesh bags were first deployed in mid-June 2014 at the center of each subplot perpendicular to the ground, where the 10 cm soil cores were taken. The mesh bag was then buried in soil. We used a total of 96 mesh bags in 2014, one for every subplot (8 subplots in 8 burned plots in addition to 8 subplots in 4 unburned plots). We harvested these mesh bags in early October 2014. In 2015, we increased the number of mesh bags to three for each subplot, totaling to 288 bags. We deployed the mesh bags in mid-June 2015. These mesh bags were harvested in early October 2015.

Each year, the mesh bags were transported to the laboratory for extraction after harvesting. We rinsed the sand and mesh with deionized water. Hyphal biomass was collected and weighed. The samples were freeze-dried and stored at -20°C after weighing.

### 2.8. Data analysis

To test whether our clipping treatment had significant effects on burned and unburned areas, we analyzed the effects of clipping treatment, fire management, PAR, soil water content, herbaceous cover, and percentage total cover all with respect to average RGR for each subplot. Hyphal biomass was measured as an effect of clipping and fire. We used one-way Analysis of Variance (ANOVA) to test clipping and fire effects on RGR, and on hyphal biomass. When considering other continuous covariates and their interaction with clipping, we used Analysis of Covariance (ANCOVA) to test their effect on RGR.

## 3. Results

### 3.1. Fire effects

To determine whether a low-intensity fire regime increased the growth of white oak seedlings, we compared the effect of fire management on RGR of each of our planted trees. Fire-managed plots, on average, had 54% higher RGRs than ones that were unburned (F (1,46) = 0.483, p = 0.511). Figure 2 shows burned plots having higher RGRs and a smaller range of rates than unburned plots.

**Figure 2:**
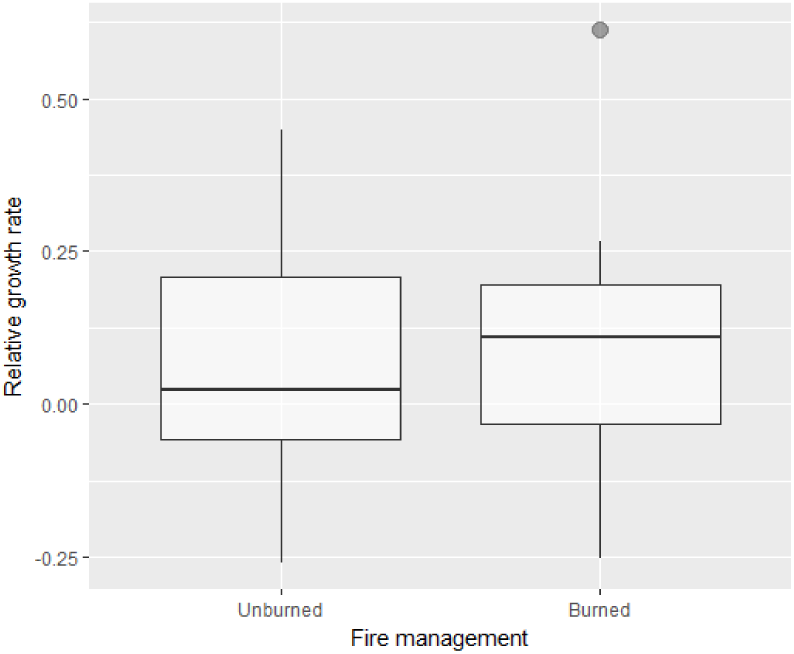
Box and whisker plot of relative growth rate for burned and unburned sites. Relative growth rate in burned sites contains an outlier represented by the closed dot.

We also tested the effect of fire management on herb cover amounts from 2015. There is a significant positive effect of fire management on herbaceous cover with burned plots having 135% more herbs than unburned plots on average (F (1,94) = 14.227, p *<* 0.001). This indicates the woody-to-herbaceous effect of fire is present on our sites.

### 3.2. Competition for light and water

In a fire-managed forest, to test whether the removal of herbaceous plants decreased tree growth due to increased competition for light and water, we investigated the effect of clipping and control treatment on RGR of trees planted only in burned areas, controlling for PAR and soil water content. In fire-managed plots, clipping had a negative effect on RGR after controlling for PAR and soil water content (F (3, 178) = 2.478, p = 0.062).

Figure 3: Left shows on average, trees in clipped plots having lower RGRs for much of the range (approximately the lower 85%) of soil water content observed. Figure 3: Right shows that trees in clipped plots having lower RGRs under the spectrum of PAR values measured. The removal of herbaceous plants with the effects of light and water overall had a negative effect on RGR of seedlings in burned areas.

**Figure 3:**
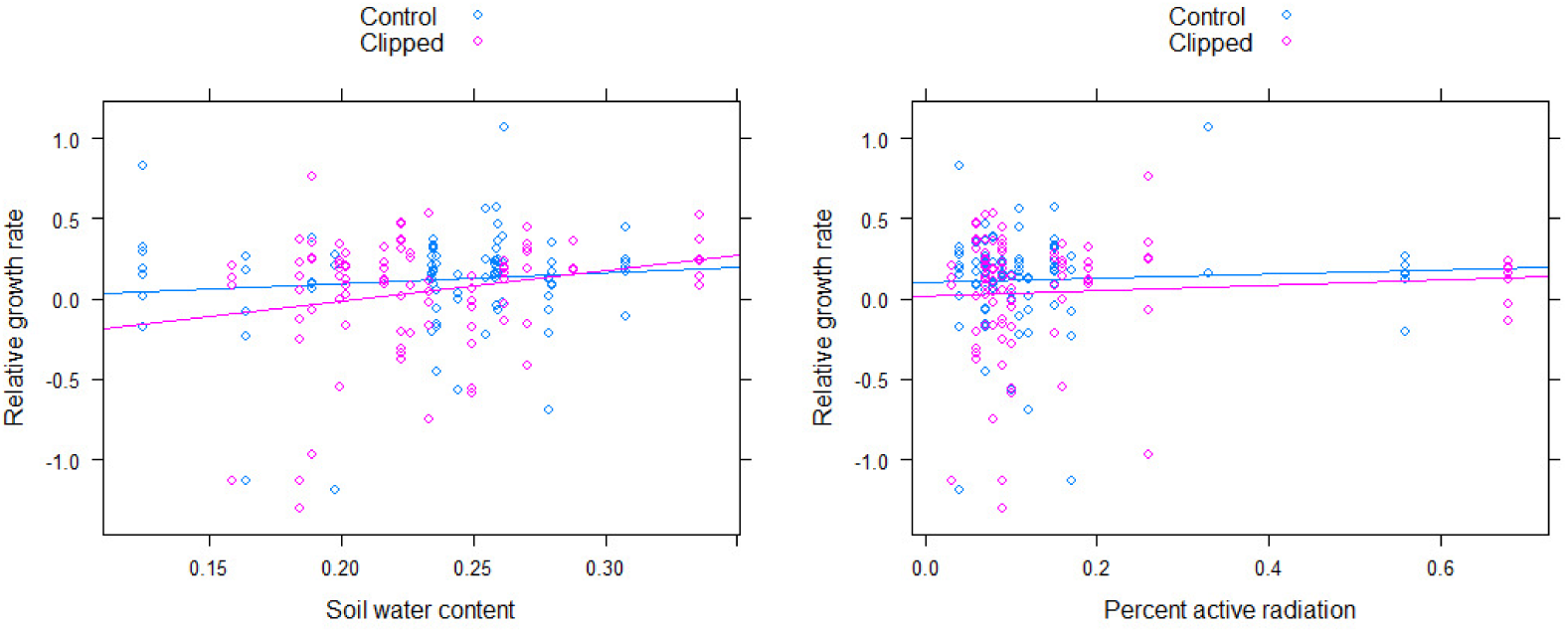
Left: Relative growth rate of white oak in burned plots as an effect of soil water content and clipping treatment. Right: Relative growth rate of white oak in burned plots as an effect of percent active radiation and clipping treatment. Both: Trees that have died or are missing are omitted. Treatment: Control, n = 90. Clipped, n = 92. Solid lines are lines of best fit.

### 3.3. Facilitation

Similarly, to assess whether the removal of herbaceous plants decreased tree growth due to the reduced protection from environmental stressors (reduced facilitative effects) in a fire-managed forest, we tested the significance between clipping and control treatments on RGR of the trees in burned areas, controlling for average cover. In fire-managed plots, clipping had a significant negative effect on RGR after controlling for total cover (F (2,179) = 4.153, p = 0.017). Figure 4: Left shows that in areas with higher percent cover (approximately the upper 75% of the observed data), trees on clipped plots had lower RGRs.

**Figure 4:**
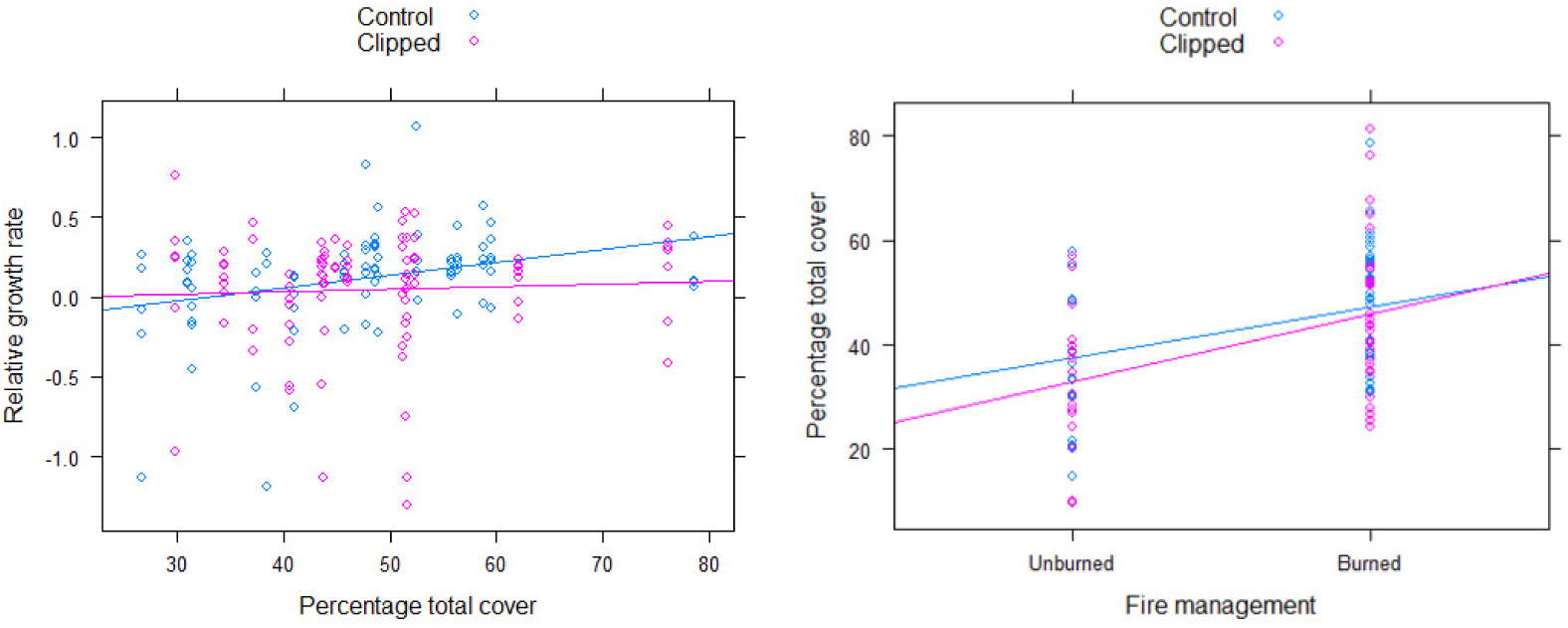
Left: Relative growth rate of white oak in burned plots as an effect of percentage total cover and clipping treatment. Trees that have died or are missing are omitted. Treatment: Control, n = 90. Clipped, n = 92. Right: Percentage total cover in all subplots as an effect of fire management and clipping treatment. Treatment: Control, n = 48. Clipped, n = 48. Fire management: Unburned, n = 32. Burned, n = 64. Both: Solid lines are lines of best fit.

In addition to the negative effect on RGR of seedlings in fire managed plots, we performed a supplementary ANCOVA to test the significance of clipping across burned and unburned areas. Clipping treatment had a significant effect of removing cover controlling for fire management (F (3, 92) = 5.361, p = 0.001). Figure 4: Right shows lower percentage total cover in clipped plots for both management areas.

### 3.4. Mycorrhizae

To determine if a mycorrhizal network increased as a result of induced herbaceous plant stress, we evaluated the effects of clipping and control treatment on hyphal biomass. Clipping treatment had a negative effect on hyphal biomass (F = (1,44) = 2.92, p = 0.094). Figure 5 shows clipping treatment correlating to lower hyphal biomass.

**Figure 5:**
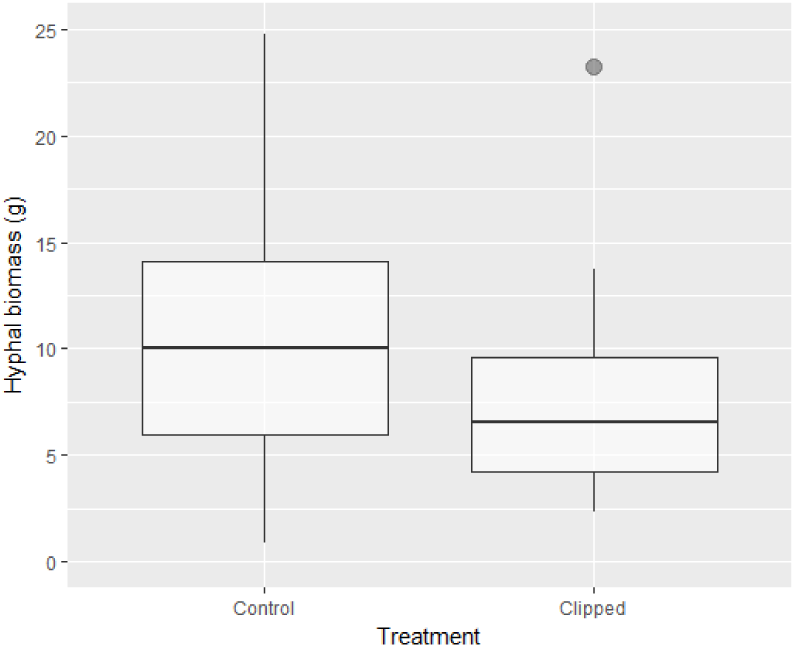
Box and whisker plot of hyphal biomass for control and clipped treatment. Hyphal biomass in clipped plots contains an outlier represented by the closed dot.

## 4. Discussion

Managers of ecosystems have often focused on restoring landscapes to healthier, more stable states where a multitude of species can thrive. Disturbance regimes, such as fire, usher in secondary succession. In an eastern deciduous hardwood forest, fire is responsible for the transition from a woody-to-herbaceous dominated understory (Peterson and Reich 2001, Knapp et al. 2015). Thus, trees and its herbaceous neighbors are both able to establish after a prescribed burn. The relationship between tree and herb species remains unclear despite widespread use of fire in woodlands. Isolating herbaceous species effects on tree fitness reveals more about their complex competitive and facilitative relationship and may improve management practices in forest ecosystems.

This experiment sought to manipulate herbaceous cover in both burned and unburned areas to observe the effects on white oak seedling fitness in select plots. We identified that compared to unburned plots, fire-managed plots yielded better performing trees. Unclipped, fire-managed plots returned the best-performing trees overall. No significant evidence of competition for water or light was found in our analyses. However, we found some support that herbaceous neighbors protected the seedlings against environmental stressors, as shown by lower RGRs after clipping was administered. This data is consistent with previous studies that support the functional use of diversity in a plant community, owing its facilitative effects to the persistence of a humid microclimate and protection against environmental severity (Wright et al. 2013, 2014). We concede that more indicators of microclimatic changes, such as humidity measurements, would better indicate if our trees were indeed protected against environmental stressors. Nevertheless, our data supports facilitative effects are at least present between seedlings and neighboring herbs.

The importance of differentiating between EM and AM was not paramount for this study, but we recognized the value of having a belowground indicator of productivity for our trees and herbs. At the root level, white oak is an EM-obligate species, and most herbs are AM-obligate (Wang and Qiu 2006). The production of mycorrhizae (specifically AM) was previously examined to increase with abiotic stress (Latef et al. 2016). We did not distinguish clipping as an abiotic or biotic stressor to our herbs. Regardless, clipping is a biotic stressor due to the alteration of plant physiology. We found hyphal biomass to correlate negatively with clipping treatment, which does not support our hypothesis. In light of this, we speculate that due to direct physical changes to the plant, some clipped herbs may have died, thus nullifying the need for any symbiotic fungi. Our findings do not construe that a dual EM-AM environment would benefit white oak seedlings. It would be problematic to assume that in our study, plot-level hyphal biomass measurements would be an appropriate determinant of individual tree fitness. Despite this, we acknowledge other studies have uncovered some benefits of a dual EM-AM environment for other oak species early in their ontogeny (Dickie et al. 2001, Egerton-Warburton and Allen 2001).

Additionally, we remark the significance of soil moisture in our study. Using Akaike information criterion corrected for finite samples (AICc), we found soil moisture to be the simplest and best indicator of RGR in all our trees (Table 1). We compared the p-values of models fitting RGR with fire management, clipping, fire-management interacting with clipping, and soil water content, and found the interaction of RGR and soil water content to be significant (p = 0.035). This result is not unexpected, as legacy studies, as well as modern experiments in different biomes, have heralded soil moisture as the catalyst to tree growth (Bassett 1964, Borchert 1994, Kraaij and Ward 2006).

**Table 1:** Model comparison among four linear models showing the effects of fire-management (mgmt), clipping, the interaction of fire management and clipping, and soil water content (swc) on relative growth rate (RGR). The RGR ~ swc model is bolded to signify a p-value < 0.05.

| Model | Coefficient | Std. Error | p-value | AICc |
| --- | --- | --- | --- | --- |
| RGR ∼ mgmt | 0.03845 | 0.05808 | 0.511 | -18.86128 |
| RGR ∼ clipping | -0.03617 | 0.05481 | 0.512 | -18.85828 |
| RGR ∼ mgmt * clipping | -0.09956 | 0.11737 | 0.672 | -15.18704 |
| **RGR ∼ swc** | 1.4798 | 0.6832 | **0.035** | -23.0678 |

Our study encountered several limitations when analyzing data. We did not conduct a variety of field tests that could have further supported facilitation in our analysis. Earlier, we mentioned humidity as a possible microclimate metric to measure. There is value in including wind speed and direction, precipitation, and longer-term measures of PAR to better gauge environmental severity in the plant microclimate. Keeping with our assumptions about labeling such relationships simply as competitive or facilitative, we may be incorrectly implying that the relationship would persist throughout the lives of the tree and herbs. Notwithstanding, facilitative dynamics in this study are only applicable to the growing season. These limitations are subject to setbacks when attempting to extrapolate to longer temporal scales, so we recommend taking year-long measurements to better understand seasonal fluctuations. Moreover, our description of facilitation is rather one-way in this experiment because of our narrow focus on positive effects for the white oak.

Future studies would ideally focus on extended and deeper observations of the facilitative dynamic. Data on actual colonization rates of mycorrhizae on individual plants under stress would be of great benefit. Because this study only focused on discrete homogeneously sized parcels of land, we have a bounded image of how the aboveground and upper parts of belowground look under prescribed burn and clipping treatment. With individual trees affected by varying levels of environmental severity, there is a need to carefully examine tree life-history strategies and their resilience in the absence of facilitative neighbors. Studies may examine the effects of trees on grass systems to resolve the one-way facilitation presumption. Looking at systems that are high in biodiversity may explain facilitation more comprehensively than interactions between a couple of grass species to a single tree species, as observed in this study. Furthermore, it may be beneficial to observe the ever-changing complexity of the mycorrhizal network in the absence of fire management, as it transitions to a predominantly woody ecosystem.

## 5. Conclusion

We have provided some evidence that facilitation is present between trees and herbaceous vegetation. The interaction of fire management and clipping treatment positively affected the RGRs of our white oak seedlings. Removal of neighboring herbs did not show higher growth rates for our trees, thus not supporting competition in our experiment. The substantial amount of biomass removed due to clipping treatment may have weakened plant communities’ defenses to environmental stressors in addition to altering the plant microclimate. Belowground, the mycorrhizal network decreased as a result of herbaceous clipping. As more forests begin or continue to be managed by fire, it is imperative to know if a facilitative or competitive dynamic is present for growing and recruiting trees. Understanding how plant communities function will shape what management practices will be developed and employed in all forest types.

## Acknowledgements

This study was supported by the University of Illinois College of ACES Office of Research. We thank our colleagues and professors from the University of Illinois who provided insight and expertise that greatly assisted the research. Thank you to Valarie Repp and Jennifer Woodyard for their immense contributions in the second year of this study. Thank you to Dr. Jennifer Fraterrigo for providing lab space and storage. Thank you to Jim Kirkland for helping us during the field seasons. Special thanks goes to Tyler Refsland for conceiving the project and guiding the study.

